# Myosin modulator Aficamten inhibits force in cardiac muscle by altering myosin’s biochemical activity without changing thick filament structure

**DOI:** 10.1101/2025.05.14.654110

**Authors:** Saffie Mohran, Kristina B. Kooiker, Ateeqa Naim, Matvey Pilagov, Anthony Asencio, Kyrah L. Turner, Weikang Ma, Galina Flint, Siyao Jiang, Jing Zhao, Timothy S McMillen, Christian Mandrycky, Max Mahoney-Schaefer, Thomas C. Irving, Bertrand C.W. Tanner, Neil M Kad, Michael Regnier, Farid Moussavi-Harami

**Author notes:** Correspondence to: Michael Regnier or Farid Moussavi-Harami, 850 Republican Street, D353, Seattle WA 98109. Authors contributed equally (Saffie Mohran, Kristina B. Kooiker).

## Abstract

**Background:** Inhibiting contractility by targeting cardiac myosin is an effective treatment for patients with hypertrophic cardiomyopathy (HCM). Aficamten is a second in class myosin inhibitor with promising clinical data showing improvements in hemodynamics and symptoms in patients with HCM. While it is known that Aficamten inhibits force and cardiomyocyte contractility by stabilizing the weak pre-powerstroke conformation, effects on myosin structure and kinetics during loaded contraction are lacking.

**Methods:** Permeabilized porcine cardiac tissue and myofibrils were used for single-molecule imaging of ATP turn over, X-ray diffraction, and mechanical measurements. Engineered heart tissues from human induced pluripotent stem cell cardiomyocytes were used to evaluate effects on force and contraction kinetics.

**Results:** In contrast to Mavacamten, Aficamten does not structurally sequester myosin heads along the thick filament. Aficamten inhibits ATPase activity by shifting myosin heads from higher to slower ATPase state, with the emergence of a super slow biochemical nucleotide turnover state. This results in decreased force and calcium sensitivity without altering cross-bridge cycling. These contractile mechanical changes are comparable to Mavacamten. Our myofibril mechanical assay showed inhibition of force with accelerated relaxation. In EHTs, while Mavacamten and Aficamten inhibit cardiac twitch forces, Mavacamten reduces the activation kinetics while both result in faster relaxation.

**Conclusions:** We used a combination of biochemical and biomechanical assays to show that Aficamten inhibits myosin ATPase without appreciably altering myosin structure. This is different from Mavacamten that strongly affects both. While both compounds inhibit contractility, differences in mechanisms of action and kinetics of force activation and relaxation could allow use in different patient populations.

## Introduction

Hypertrophic cardiomyopathy (HCM), the most common genetic heart disease with prevalence of around 1:500, is defined by left ventricular hypertrophy that is not attributed to hypertrophy from another cardiac, metabolic or systemic process. It can be accompanied by hyperdynamic contraction, outflow tract obstruction, and impaired ventricular relaxation.^1^ A genetic cause is identified in about 30-60% of patients diagnosed with HCM and variants in genes encoding for sarcomeric proteins are among the leading causes.^2^ Mavacamten (Mava) was recently approved for use in patients with obstructive HCM as it improved exercise capacity and left ventricular outflow track obstruction.^3,4^ Mava selectively binds to myosin and inhibits cardiomyocyte contractility.^5,6^ Studies into the mechanism of Mava have demonstrated that it decreases the rate of inorganic phosphate (Pi) release and stabilizes the OFF state of myosin on the thick filament.^7–10^ In addition to Mava, a second small molecule myosin inhibitor, Aficamten (Afi), recently completed a phase III clinical trial that showed improvement in symptoms and functional capacity in patients with obstructive HCM.^11^

Despite being discovered using a similar myofibril ATPase assay, Afi has a distinct binding site and mechanism of action that differs from Mava. A recent study showed that Afi and blebbistatin (Bleb) share a common binding site on cardiac myosin that is distinct from Mava.^12^ The study used saturating concentrations in biochemical assays, to investigate how Afi impacts myosin function. Their key conclusions suggest that Afi reduces the rate of actin-activated Pi release and modestly decreases actin-activated ADP release from cardiac myosin S1. They also suggest that Afi stabilizes the weak pre-powerstroke conformation (ADP.Pi), resulting in a loss of force bearing myosin heads and a reduction in contractility.

Here, we provide a comprehensive mechanistic study on how Afi impacts the structural, biochemical, and contractile function of myosin within the native sarcomeric structure. By utilizing porcine and human induced pluripotent stem cell (iPSC) models for our assays, we observe how Afi impacts cardiac β-myosin cross-bridge recruitment and cycling under physiological conditions.

## Materials and Methods

The data that support the findings of this study are available from the corresponding author upon reasonable request.

### Animal use and ethics

All experiments followed protocols approved by both the University of Washington and the Illinois Institute of Technology Institutional Animal Care and Use Committees according to the “Guide for the Care and Use of Laboratory Animals” (National Research Council, 2011). Muscle tissue was collected in accordance with the U.K. Animals (Scientific Procedures) Act 1986 and associated guidelines. Farm pig hearts or Yucatan mini pigs (Exemplar Genetics) were obtained immediately after the animal was euthanized and rinsed in cold oxygenated Tyrode’s buffer.

### Myofibril single-molecule imaging of ATP Turnover

Myofibril isolation and flowcell preparation can be found in the supplemental methods section. Flowcells were prepared as described previously^13^ and connected to a syringe pump (WPI, AL-1000) at one end and at the other, to a 1.5 ml microcentrifuge tube with holes to allow access of pipette tips. 200 µl of prepared porcine myofibril suspension (supplemental methods) was added to the 1.5 ml microcentrifuge tube and drawn into the flowcell at 1 ml/min then washed back through at 0.2 ml/min. To allow adhesion of myofibrils to the PLL-coated surface, flowcells were incubated on ice for 30 minutes. Excess non-adherent myofibrils were withdrawn from the flowcell using 600 µl Prep buffer (supplemental methods) at 10 ml/min.

Myofibrils were incubated with Cy3-ATP buffer (Prep buffer + 1 µM Cy3-ATP, 500 µM ATP, 0.5 mM phosphoenolpyruvate (PEP), 2.2 units Pyruvate kinase (PK)) for 30 mins at RT. Where used, Aficamten (MedChemExpress, Cat. No.: HY-139465) solubilized in 100% DMSO, was added to the Cy3-ATP buffer and Chase buffer (Prep buffer + 5 mM ATP, 5 mM PEP, 22 units PK) at final concentrations stated in Figure 1. (Final DMSO remained constant at 0.06% v/v). Following this, a single frame was taken of the myofibril fully saturated with Cy3-ATP, followed by a rapid wash with Chase buffer at 10 ml/min.

**Figure 1:**
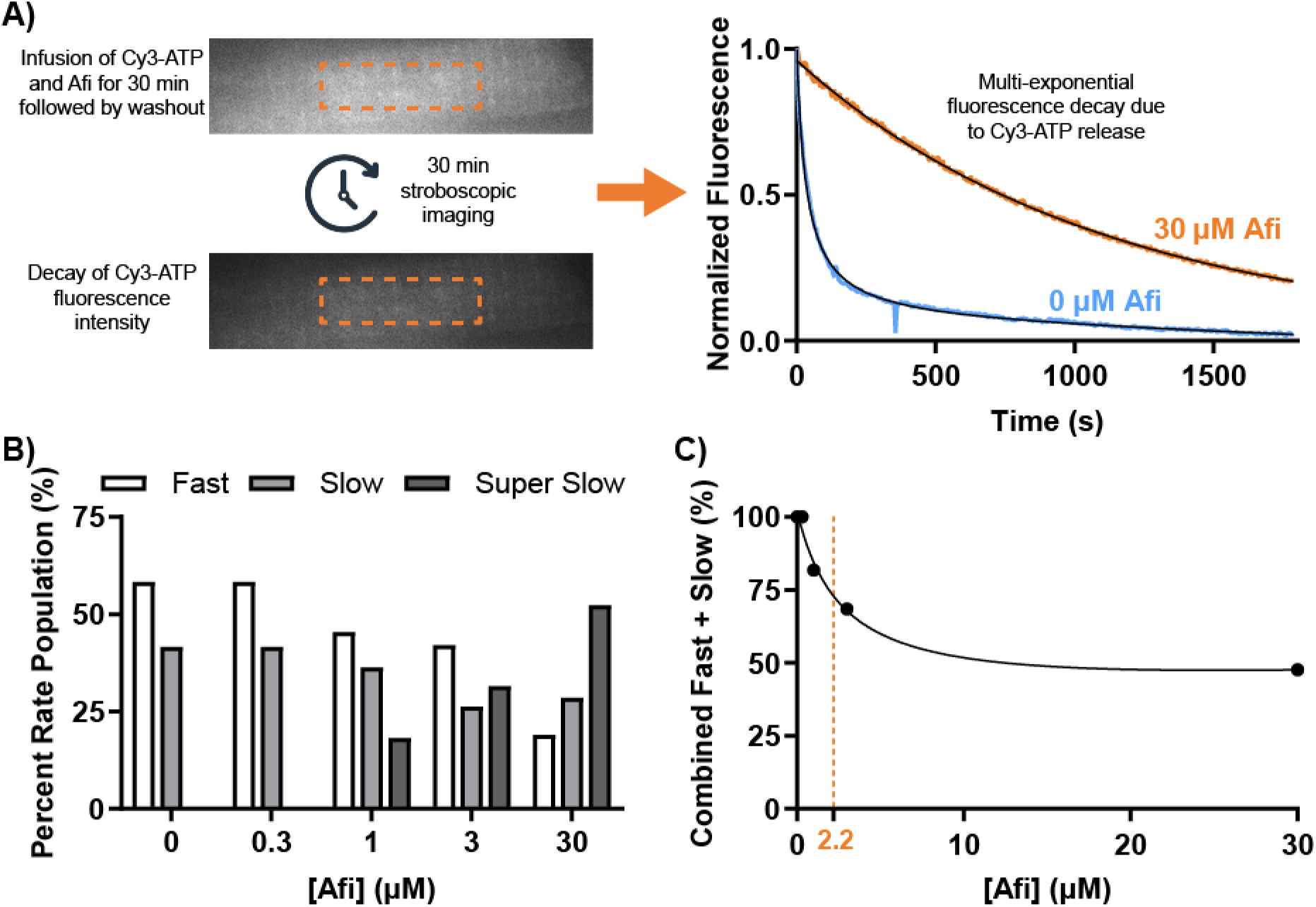
Increasing concentration of Afi results in the emergence of a very slow ATPase state. A) Experimental protocol used, showing region analyzed (in box) for single-molecule ATP imaging (left) and example tracing of normalized fluorescence in absence (blue) or presence (orange) of 30 μm Aficamten (right). B) The percentage of all rate constants in the fast, slow, or super slow state at each concentration of Afi shows the emergence of the super slow state at 1 μM Afi and above. C) To calculate the K_d_ of 2.21 (± 0.76) μM (orange dashed line), all the fast and slow state percentages at each concentration were combined and fit to a weak binding equation (S/(K+S)). N = 6-15 myofibrils were measured at each concentration of Afi then rate constants were binned into fast (0.05 - 0.005 s^−1^), slow (0.005 - 0.0005 s^−1^), or super slow (< 0.0005 s^−1^).

To increase efficiency, flowcells were mounted onto a custom-built automated stage programmed to move between user-specified positions along one axis. Between each frame images from neighboring myofibrils were taken. Use of this automated stage allowed up to 4 myofibrils to be simultaneously imaged during the 30-minute imaging period.

All imaging was carried out at room temperature (RT) using a custom-built oblique angle fluorescent (OAF) microscope.^14^ The sample was illuminated for 15 ms every 5 seconds using a 561 nm laser (OBIS LS laser, Coherent, USA) to excite Cy3-ATP, for a total of 30 mins.

To extract the rate of fluorescence decay for each myofibril, an ROI was drawn using ImageJ to encompass the entire myofibril and the total intensity profile was extracted over time. These data were transferred to Excel and then fitted to a triple exponential decay, providing both rate constant and amplitudes.

### X-ray diffraction

Wild type Yucatan mini-pig hearts were provided by Exemplar Genetics LLC. Permeabilized tissue preparation and beamline specifications are described in as previously described.^15^ X-ray diffraction experiments were performed at the BioCAT beamline 18ID at the Advanced Photon Source, Argonne National Laboratory.^16^ Skinned muscle preparations were mounted in a custom rig. The muscle was incubated in a customized chamber with the solution temperature at 28 °C. The SL of the muscles was set to 2.3 µm by measuring the diffraction pattern of the fiber utilizing a helium-neon laser (633 nm). X-ray fiber diffraction patterns were collected in pCa 8.0 solution in the absence or presence of 50 μM of Afi. Fibers were incubated for 15 minutes in 50 µM Afi prior to imaging. Analysis of x-ray diffraction images were done as described previously^17,18^.

### Demembranated tissue mechanics

Tissue preparation and solutions are described in supplemental methods.

#### Isometric force and rate of tension redevelopment

Porcine left ventricular (LV) tissue was demembranated overnight at 4 °C in a 50% glycerol relaxing solution with 1% Triton X-100. Permeabilized LV strips were dissected to ∼120 x 600 μm and mounted between a force transducer (Aurora Scientific, model 400A) and a motor (Aurora Scientific, model 315C) using aluminum T-clips (Aurora Scientific).^19^ The tissue was stretched to a sarcomere length (SL) of ∼2.3 μm for each experiment. Experiments began in a relaxing solution with pCa (= −log_10_[Ca^2+^]) of 8.0, then moved through a pCa curve from 6.0 to 4.5 in physiological solution (pH 7.0) at 21 °C with 3% dextran. Experiments were done in three groups of untreated (ND), 1 µM Afi, or 1 µM Mava (MedChemExpress). The tissue was incubated in relaxing conditions with or without compound for 2 hours at room temperature (RT) prior to experimentation. During experimentation, tissues were allowed to reach stead-state force (F) at each pCa. F-pCa curves were collected with custom code using LabView software and fit to the Hill equation in GraphPad Prism. The rate of tension redevelopment (*k*_TR_) was measured at each pCa with a 15% slack and re-stretch.

#### Sinusoidal length-perturbation analysis of viscous modulus

Permeabilized porcine myocardial strips (∼180×700 μm) were mounted between a piezoelectric motor (P841.40, Physik Instrumente, Auburn, MA) and a strain gauge (AE801, Kronex, Walnut Creek, CA) using aluminum T-clips. Each strip was lowered into a 30 μL droplet of relaxing solution (pCa 8.0) maintained at 28 °C. SL was then set to 2.3 μm, and the solution was exchanged with activating solution (pCa 4.8). Sinusoidal length perturbations of 0.125% myocardial strip length (clip-to-clip) were applied at multiple frequencies from 0.125 to 250 Hz to measure the complex modulus as a function of frequency.^20^ The complex modulus describes viscoelastic myocardial stiffness which arises from the change in stress (=force per cross sectional area) divided by the change in muscle length that is in-phase (=elastic modulus) and out-of-phase (=viscous modulus) with the oscillatory length change at each discrete frequency.

The viscous modulus at pCa 4.8 was fit to a polynomial to extract the maxima and minima frequencies for each condition, where the “dip frequency” describes force-generating events and cross-bridge recruitment while the “peak frequency” describes cross-bridge distortion events and cross-bridge detachment rate.^21–23^ These characteristic “dip” and “peak” regions of the viscous modulus were used to assess the effects of Dani and Mava on cross-bridge cycling kinetics under maximal Ca^2+^-activated conditions.

### Myofibril mechanics

Myofibril activation and relaxation measurements were performed on a custom setup as previously described.^24,25^ Myofibrils were isolated from LV porcine tissue using a tissue homogenizer for 10 s. Myofibrils were then platted onto the custom setup were they were mounted between a glass force transducer and an inflexible motor arm. A dual photodiode system measured myofibril force by recording needle deflection. The force transducer needle stiffness measured 7.98 µm/µN. A double-barreled glass pipette delivered relaxing (pCa = 8.0) and activating (4.5) solutions to the isolated myofibril using a rapid switching technique. Activation and relaxation data were collected at 21 °C and fitted as previously described.^25^

### Human induced pluripotent stem cell differentiation and engineered heart tissue casting

Differentiation of WTC11 human induced pluripotent stem cells (hiPSC) into cardiomyocytes (CM) is detailed in the supplemental methods. To cast hiPSC-CMs into engineered heart tissues (EHTs), 500k 21 days old lactate-purified WTC11 hiPSC-CMs were mixed with 50k HS27a (human marrow stromal cells^26^), 5 mg/ml fibrinogen (Sigma), and 3 unit/ml thrombin (Sigma) then cast onto wells with PDMS post arrays^27^. Gelation occurred over the following hour and a half, then tissues were moved to a fresh 24 well plate with RPMI + B27 + insulin + 5 g/L aminocaproic acid (Sigma) and stored at 37°C. Media was changed every 2-3 days while the tissue compacted and matured.

### Intact twitch assay

EHTs were cut off posts 2 weeks after casting (35 days post-inducement of CM differentiation) and moved into a dish containing DMEM/F12 (Gibco) continuously bubbled with 95% O_2_/ 5% CO_2_ to maintain physiological pH and mounted between two platinum omega clips. The EHT was then mounted into the Intact Muscle Chamber System (IonOptix) and perfused with DMEM/F12 supplemented with CaCl_2_ (final Ca^2+^ = 1.8 mM) continuously bubbled with 95% O_2_/ 5% CO_2_. EHTs were electrically paced at 1 Hz and maintained at ∼32 °C throughout the experiment. EHTs were given 15 minutes to stabilize, during which time optimal length was set to where the total amplitude of the twitch no longer increased. Drug titrations were performed with 0.01 % DMSO (ND), then 1 and 2 μM Afi or 0.5 and 1 μM Mava with 15 mins incubations at each concentration. For each condition, 30 second traces were recorded and analyzed using IonWizard software (IonOptix).

### Statistical Analysis

We used GraphPad Prism 10 for data presentation and statistical analysis unless otherwise stated. The data is presented as mean ± standard error of the mean. For porcine isometric demembranated tissue mechanics and myofibrils, we used an Ordinary one-way ANOVA with Tukey’s multiple comparisons test. For viscous modulus measurements, SPSS software (IBM Statistics, Chicago, IL) was used. Nested linear mixed models were utilized for the minimum or maximum frequency parameters determined from polynomial curves fit from the viscous modulus. Post hoc analysis used Fisher’s least significant difference test. For X-ray diffraction and EHT twitch analysis, we used paired two-tailed t-tests. For *in vitro* motility, we used two-way ANOVA with Šídák’s multiple comparisons.

## Results

### Aficamten inhibits ATPase activity without structurally sequestering myosin heads on the thick filament

To understand the effect of Afi on ATPase activity, we assessed the decay of fluorescently labeled ATP (Cy3-ATP) in permeabilized porcine cardiac myofibrils as myosin cycles and releases Cy3-ADP (Figure 1A). Experiments were performed on relaxed myofibrils after infusion with Cy3-ATP and Afi at concentrations ranging from 0 to 30 μM for 30 minutes. This was followed by a wash-out of Cy3-ATP with unlabeled ATP (with Afi) and 30 minutes of stroboscopic imaging to image the decay in fluorescence intensity. The intensity was then fit to a multi-exponential decay to extract multiple rate constants and their relative amplitudes (Figure 1A). For each myofibril, the resultant rate constants were classified as one of three ATP turnover states (fast, slow, super slow). Figures 1B and S1 show that as the concentration of Afi increases, the population of myosin heads in the fast state decreases, with the emergence of a super slow state at 1 μM Afi. Interestingly, the population in the slow state seems to only minimally decrease with increasing Afi. To plot a binding isotherm for Afi, we combined the fast and slow states, which represents myosin heads not in the super slow ATPase state and fit a weak binding equation (S/(K+S)) to calculate a K_d_ of 2.21 (±0.76) μM (Figure 1C). This is in the same range as was seen by Hartman and colleagues^28^.

In addition to assessing the biochemical inhibition of myosin ATPase activity by Afi in permeabilized porcine tissue, we utilized small angle x-ray diffraction to assess the impact on thick filament structure. Previous studies have shown that myosin modulators can significantly impact the position of myosin and the organization of the thick filament structure.^15,18^ Paired x-ray diffraction images were collected under relaxed conditions (pCa 8.0) with no drug (ND) and in the presence of 50 μM Afi (Figure 2A). Measures of the equatorial reflections showed no significant difference in the lattice spacing (d_1,0_; FigS2) or intensity ratio (I_1,1_/I_1,0_; Figure 2B). Quantification of the radius of the center of mass of the myosin heads (R_m_; Figure 2C), which describes the average distance of the myosin heads from the thick filament, also showed no significant difference in the presence of Afi. These results show that in saturating conditions, Afi does not induce changes to the sarcomere inter-filament spacing or the position of the myosin heads relative to the thick and thin filaments. Additional assessment of the axial spacing from the third-order myosin-based meridional reflection (S_M3_; Figure 2D) and the sixth-order myosin-based meridional reflection (S_M6_; Figure 2E) showed no change in the presence of Afi. These spacing measurements indicate that Afi does not induce any structural change in position between the myosin crowns (S_M3_) or alter the thick filament backbone (S_M6_) in a manner typically associated with strain and mechano-sensing.

**Figure 2:**
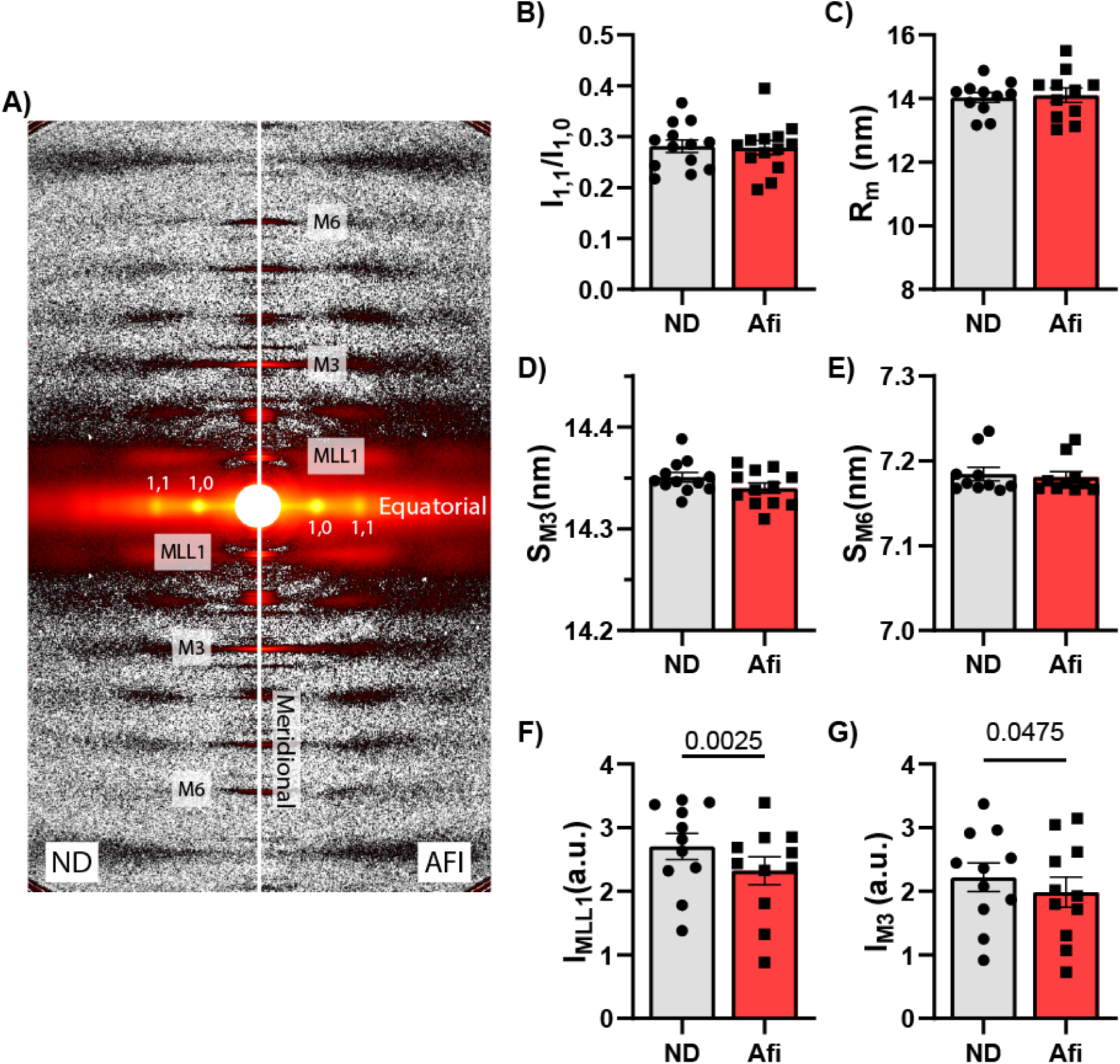
50 μM Afi minimally effects thick filament structure in relaxed porcine tissue. A) Representative X-ray diffraction patterns before (ND; left) and after (AFI; right) incubation in 50 μM Aficamten. Afi does not affect the (B) intensity ratio (I_1,1_/I_1,0_) or the (C) radius of the cross-bridge center of mass (R_m_) suggesting myosin heads are not moving toward or away from the thick filament backbone due to Afi. The spacing of the (D) third-order of myosin-based reflection (S_M3_) and (E) sixth-order of myosin-based reflection (S_M6_) are not changed. The intensity of the (F) third-order myosin-based reflection (I_M3_) and (G) first-order myosin-based layer line (I_MLL1_) slightly decrease after Afi incubation, suggesting Afi causes some disorder in the myosin heads. N = 13 tissues samples were prepared and measured. Paired t-tests were performed for each measurement.

Interesting, both the intensity of the first-order myosin-based layer line (I_MLL1_; Figure 2F) and third-order myosin based meridional reflection (I_M3_; Figure 2G) in the presence of Afi decreased by 10.5% compared to ND paired controls. The decrease in I_MLL1_ and I_M3_ describes a reduction in the number of quasi-helically ordered myosin heads on the thick filament.

Both the equatorial and meridional results with Afi are different from those described by Anderson et al and Ma et al for Mava.^7,18^ They reported that Mava significantly shifts the radial position of myosin towards the thick filament backbone and increases the helical ordering of the myosin crowns. These structural modulations caused by Mava, coupled with the ATPase inhibition, result in significantly inhibited actomyosin interaction and sarcomeric contractility.

### Aficamten inhibits contractile force without altering crossbridge cycling kinetics in permeabilized porcine tissue

To assess how the biochemical and structural effects of Afi impacted cardiac contractility, we first examined steady-state force production, calcium sensitivity, and myosin cycling kinetics in permeabilized porcine tissue. Figure 3A shows the concentration dependence of the time required to reach steady state inhibition by Afi. At higher Afi concentrations, such as those used for the ATP turnover (Fig 1) and X-ray diffraction (Fig 2) experiments, force is fully inhibited after 10 minutes. Therefore, the remaining steady-state force measurements were performed at sarcomere length of 2.3 μm after a two-hour incubation either in the absence of drug (ND; 0.1 % DMSO) or in the presence of 1 μM Afi or Mava. Afi and Mava had similar inhibitory effects for isometric contraction measurements (Fig 3B-F). Both drugs shifted the force-pCa relationship to the right (Fig 3B), with significantly decreased maximal calcium activated force (Fig 3C) and calcium sensitivity of force (pCa_50_; Fig 3D). There were no changes in the Hill coefficient (Fig 3E), suggesting no changes in cooperativity of contractile activation. There were also no changes in cross bridge cycling kinetics as determined by the rate of tension redevelopment (*k*_TR_; Fig 3F). Viscous modulus was measured in maximally activating conditions sweeping frequencies from 0.125 Hz to 100 Hz to assess steady state crossbridge attachment and detachment. The overall magnitude of the viscous modulus decreased (Fig 3G), which is consistent with a decrease in the number of strongly bound cross bridges. The horizontal positions of the minimum and maximum frequencies indicate differences in cross bridge kinetics due to cross bridge attachment and detachment rates, respectively. The dip in the modulus-frequency relationship (minimum viscous modulus) was shifted toward a lower frequency for Mava but not Afi (Fig 3H), suggesting that the rate of crossbridge attachment decreased only for Mava. No frequency shifts were observed for the peak viscous modulus values (Fig 3I), indicating no change in rate of crossbridge detachment.

**Figure 3:**
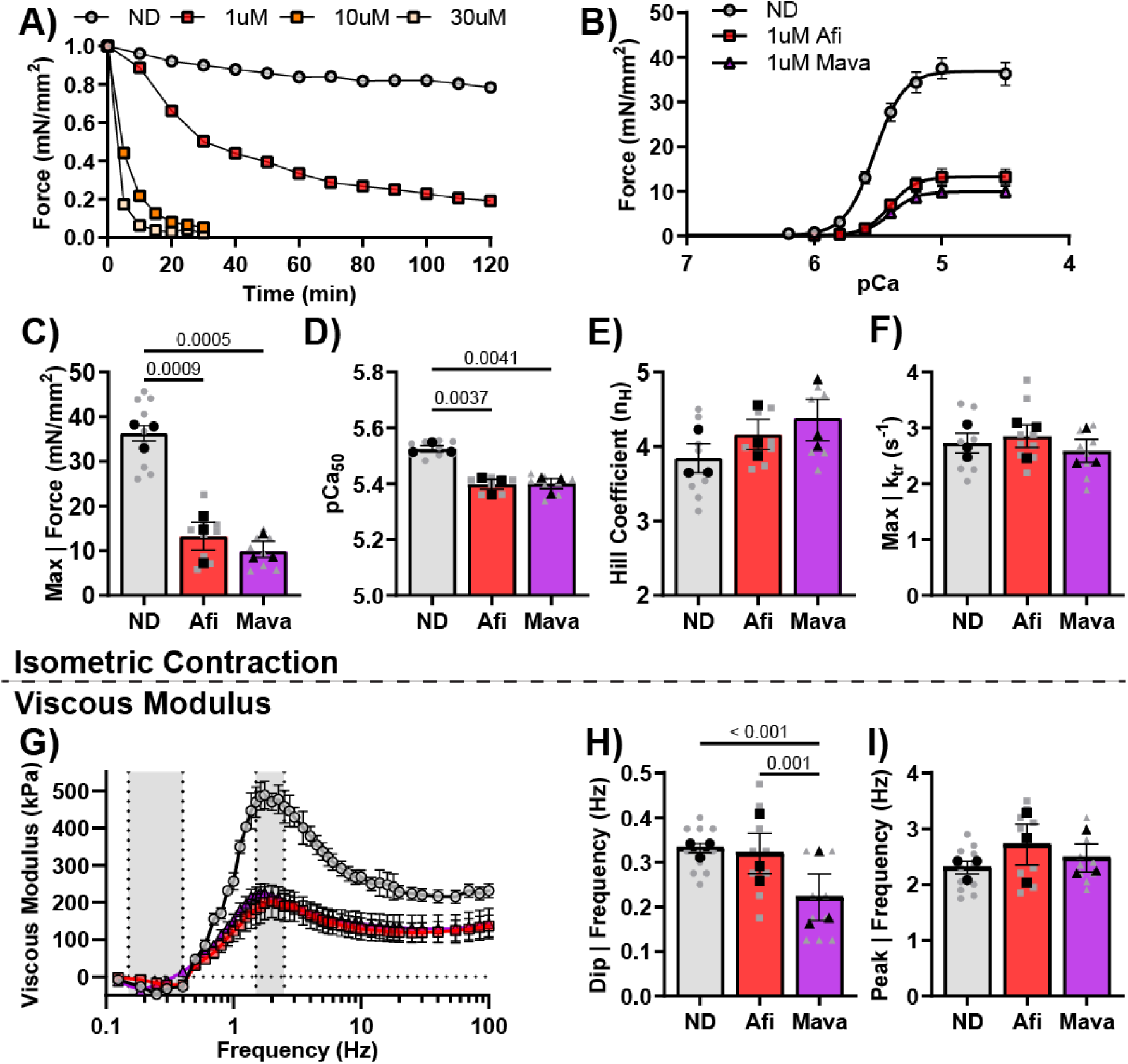
Afi and Mava inhibit force, calcium sensitivity and viscous modulus dip frequency in demembranated tissue mechanics. A) The time required for full force inhibition by Afi decreased as the concentration of Afi increased. After 2-hour incubation in ND, 1 μM Afi, or 1 μM Mava, (B) the force-pCa relationships were shifted rightward for both drugs due to decreases in (C) maximal calcium-activated force and (D) calcium sensitivity. (E) The Hill coefficient and (F) maximal calcium-activated rate constant of tension redevelopment (*k*_tr_) were not affected. (G) Viscous moduli values were plotted against frequency to assess cross-bridge binding and kinetics. At maximal activation, both drugs decreased moduli values across much of the frequency range (H) The dip frequency (minimum viscous modulus) and (I) peak frequency (maximum viscous modulus) were not affected by Afi, but the dip frequency was shifted lower for Mava suggesting slower cross-bridge attachment rate. N = 8-11 tissues samples from three different pigs were prepared and measured for each condition. Ordinary one-way ANOVAs were performed with Tukey’s multiple comparisons tests for isometric tension analysis. For viscous modulus, nested linear mixed models were utilized with post-hoc Fisher’s least significant difference test.

### Aficamten inhibits force generation and accelerates relaxation kinetics in isolated myofibril preparations

The utilization of subcellular myofibril preparations allows for the measurement of force and kinetics during the transition states between activation and relaxation. To switch the myofibril between maximal activation (pCa 4.5) and relaxation (pCa 8.0) solutions we utilize a rapid solution switching method with a double-barreled pipet. Representative traces in Figure 4A show the normalized force relationship during myofibril activation and relaxation in the presence of ND, 1.0 μM Afi, or 0.5 μM Mava.

**Figure 4:**
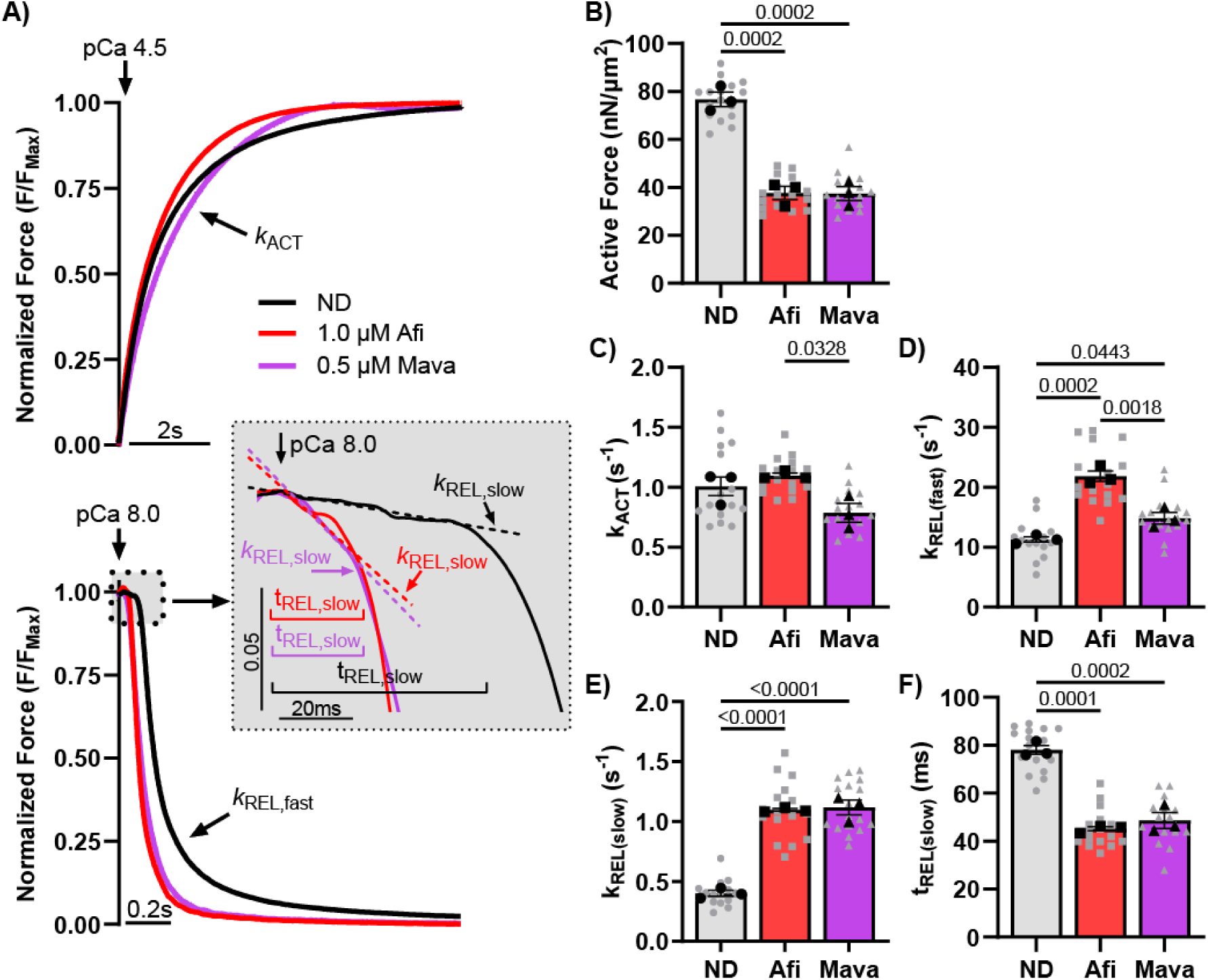
Afi and Mava inhibit maximal calcium-activated force and slow relaxation kinetics in isolated myofibrils. A) Average traces showing activation (top), the fast phase of relaxation (bottom), and the slow phase of relaxation (inset). B) Both Afi and Mava inhibited maximal calcium-activated force. (C) Neither drug changed the rate constant of activation (*k*_ACT_) versus ND. However, Afi and Mava were significantly different from each other. (D) The rate constant of the fast phase of relaxation (*k*_REL (fast)_) and (E) the rate constant of the slow phase of relaxation (*k*_REL (slow)_) were increased for both drugs versus ND, and (F) the time of slow relaxation (t_REL (slow)_) was shorter for both drugs versus ND. N = 16-17 myofibrils from three different pigs were prepared and measured for each condition. Ordinary one-way ANOVAs were performed with Tukey’s multiple comparisons tests for isometric tension analysis.

Switching between pCa 9 and pCa 4.5 did not significantly affect the exponential rate (*k*_ACT_) of force generation in the presence of either Afi or Mava (1.10 ± 0.02 and 0.79 ± 0.08 s^−1^) compared to untreated controls (1.01 ± 0.08 s^−1^) (Figure 4C). Interestingly, however, Afi and Mava results were significantly different (p=0.0328) from each other, where Afi was slightly but not significantly elevated and Mava was slightly reduced. In addition to the activation kinetics, we observed that both Afi and Mava significantly inhibited the maximally activated force (37.64±2.81 and 37.46±2.90 nN/μm^2^) of the isolated porcine cardiac myofibrils compared to untreated controls (76.70±3.00 nN/μm^2^) (Figure 4B).

Upon the rapid transition from activating to relaxing solution, we observed a biphasic relaxation relationship that consists of both an initial linear “slow phase” and exponential “fast phase” of relaxation as the force returns to baseline. The linear rate constant from the slow phase of relaxation (*k*_REL,slow_) describes the rate of cross-bridge detachment.^29^ Previous studies have shown this rate constant value is independent of the Ca^2+^ dissociation from troponin.^30^ The duration of the slow phase (t_REL,slow_), however, depends on the time of thin filament deactivation which, in turn, is dependent on the properties of thin filament regulatory proteins (troponin and tropomyosin) and the calcium concentration during activation. The final component of myofibril relaxation is the exponential rate constant (*k*_REL,fast_) that measures the rapid return of force to baseline.

This measurement characterizes multiple different inter-sarcomere dynamics and involves both active and passive properties that enable the sarcomeres to relax.^30^ Afi significantly accelerated *k*_REL,fast_ (21.9 ± 0.9 s^−1^) compared to Mava (14.8 ± 1.0 s^−1^) and to ND (11.3 ± 0.4 s^−1^, Figure 4D). Both Afi and Mava significantly increased *k*_REL,slow_ (1.10 ± 0.01 and 1.12 ± 0.06 s^−1^) compared to ND (0.40 ± 0.03 s^−1^, Figure 4E). Both compounds also significantly decrease t_REL,slow_ (45.1 ± 0.9 and 48.6 ± 3.3 ms) compared to ND (78.1 ± 1.8 ms, Figure 4F). These significant effects of Afi on myofibril relaxation kinetics were very interesting given the lack of differences in *k*_TR_ or viscous modulus frequencies.

### Aficamten decreased the tension time integral primarily through inhibition in force production in intact engineered heart tissues (EHTs)

To determine how all the reductionist preparation results integrate to alter contractile function, we compared how Afi and Mava impact twitch tension and kinetics in intact engineered heart tissues (EHTs) made from human induced pluripotent stem cell-derived cardiomyocytes (iPSC-CMs). 21-days post-onset of differentiation hiPSC-CMs were cast onto PDMS posts along with stromal cells in a fibrin gel then allowed to compact and mature for two weeks. On day 14 post-casting, EHTs were cut from the posts, secured between two omega clips, and mounted between a force transducer and post. Average twitch tension normalized to ND (0.01% DMSO) was inhibited in the presence of Afi and Mava (Fig 5A), though higher concentrations of Afi (2 μm) were required to reach similar levels of inhibition compared to Mava (1 μm, Fig 5B). The substantial decreases in force led to negative tension index values (Fig 5C), which has been shown to be predictive of hypo-contractility at the whole organ level.^19,31^ Time to peak tension (TT_P_; Fig 5D) was significantly longer for Mava but not changed for Afi. Time to 50% relaxation (RT_50_; Fig 5E) and time to 90% relaxation (RT_90_; Fig 5F) were significantly shorter for Afi, while only RT_50_ was shorter for Mava.

**Figure 5:**
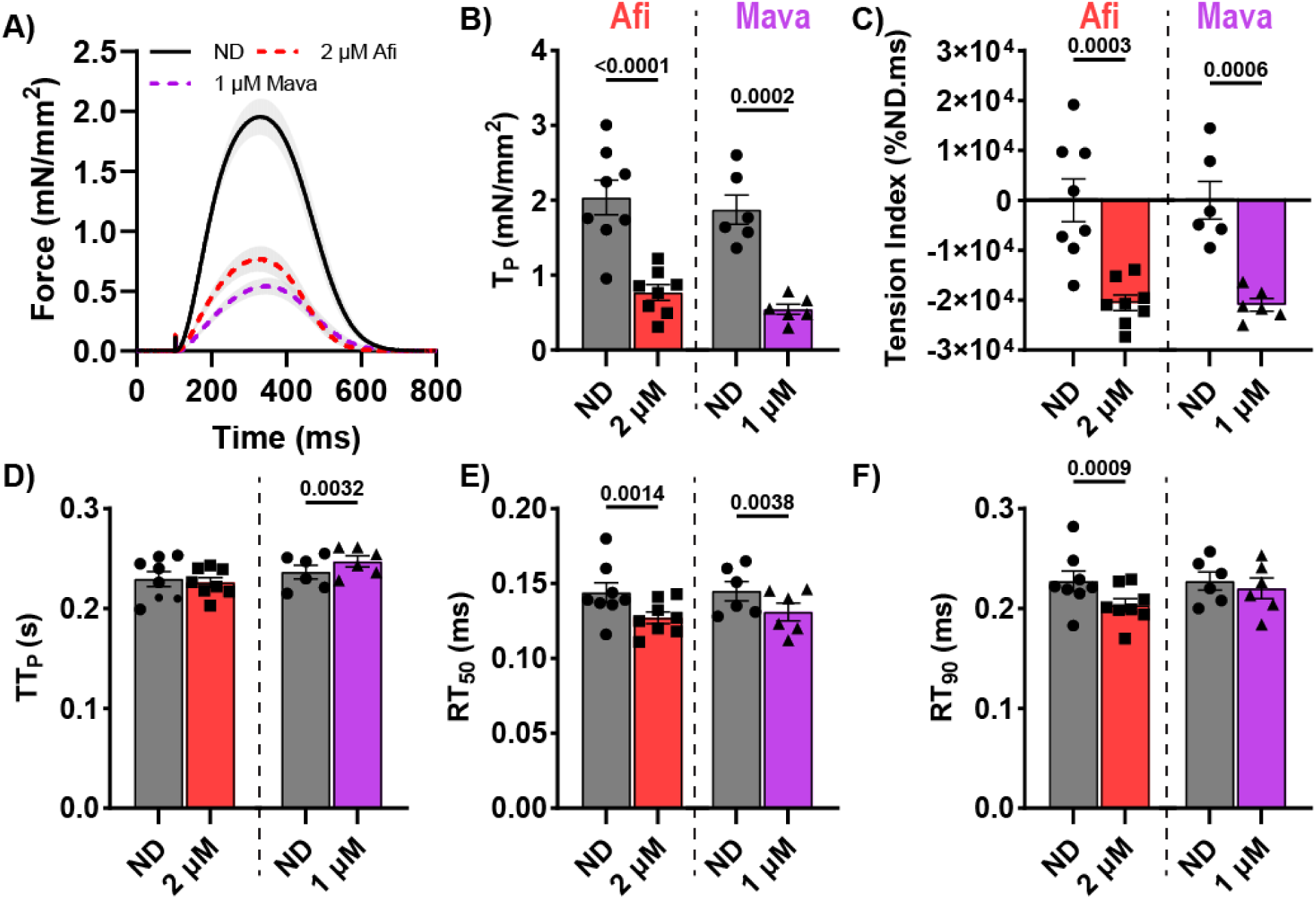
Afi and Mava decrease tension index in a concentration-dependent manner primarily through changes in peak tension in engineered heart tissues. (A) Average twitch traces of paired experiments (ND versus Afi or ND versus Mava) were normalized to EHT cross-sectional area. (B) Peak twitch tension was inhibited by both Afi and Mava leading to (C) negative tension index values for both drugs. (D) Time to peak tension (TT_P_) was longer for Mava but not Afi. (E) Time to 50% relaxation (RT_50_) was shorter for both Afi and Mava, and (F) time to 90% relaxation (RT_90_) was only shorter for Afi. N = 6 – 8 EHTs were matured and measured. Paired t-tests were performed for each EHT.

## Discussion

In this study, we investigated the biophysical and biochemical mechanisms underlying the ability of Aficamten (Afi) to reduce cardiac contractility. Afi, a second-in-class, myosin-specific small molecule currently in phase 3 clinical trials, has demonstrated functional improvements comparable to the FDA-approved Mavacamten (Mava) in patients with obstructive hypertrophic cardiomyopathy.^11,32^ Recent findings by Hartman and colleagues^28^ revealed that Afi significantly inhibits inorganic phosphate (Pi) release and may reduce ADP release rate constants in purified myosin S1 in unloaded conditions, consistent with pre-powerstroke (M.ADP.Pi) modeling. Building on these findings, we explored how Afi modulates myosin structural organization and sarcomeric contractility under load at myofibril and tissue levels.

Using a porcine cardiac model, that expresses β-myosin, we found that Afi significantly reduces the number of myosin heads available to interact with actin and produce force. Interestingly, there was an increased population of low ATPase activity myosin and emergence of a super slow ATPase activity population with increasing Afi in the absence of the structural changes monitored by x-ray diffraction. This suggests that impaired ATPase activity can be achieved without structural sequestration. Our results also suggest Afi induces significant acceleration of relaxation kinetics during myofibril measurements in saturating Ca^2+^ conditions and physiological conditions during isometric twitch measurements utilizing the engineered heart tissue platform. Our results also suggest Afi induces significant acceleration of relaxation kinetics during myofibril measurements in saturating Ca^2+^ conditions and physiological conditions during isometric twitch measurements utilizing the engineered heart tissue platform.

### The pool of recruitable myosin can be impacted by biochemical and/or structural modulation

The biochemical ATPase state (SRX and DRX) and thick filament structure (ON and OFF) have often been thought to be interchangeable. However, recent studies have provided strong evidence suggesting these biochemical and structural measurements are uncoupled under different conditions.^33,34^ Structural assessments of Mava bound to β-myosin have shown that it increases the formation of head-to-head and head-to-tail complexes in the thick filament.^35,36^ These alterations to the myosin head structure caused by increased affinity of inorganic phosphate to myosin (M.ADP.Pi) lead to increased myosin crown organization along the thick filament backbone that can be measured by structural imaging techniques such as x-ray diffraction^18^ and cryo-em.^35^ The induced head-head structural interaction also increases the inhibitory effect of Mava as shown by the significant decrease in K_i_ between isolated β-myosin S1 and HMM protein preparations (K_i_, 1.76 μM and 0.32 μM respectively)^33^.

In contrast to Mava, Afi does not induce changes in the myosin crown organization or position with respect to the thick filament, suggesting that Afi achieves ATPase inhibition through alternative mechanisms. Molecular modeling and compound competition experiments^12^ show that Afi binds in the same region of myosin as blebbistatin and requires the same M.ATP state to bind.^37^ This binding state complicates performing reductionistic assays to assess actomyosin affinity and ADP release kinetics as these require rigor (A.M.) conditions prior to mixing with nucleotide. Utilization of the modified Cy3-ATP nucleotide replacement assay enabled us to introduce Afi to myosin in the presence of ATP, enabling the compound to bind prior to performing nucleotide replacement assays. This technique allows us to perform compound titration curves and validated the emergence of the super slow state, a compound induced nucleotide replacement rate 10x slower than the described 0.002 s^−1^ SRX state. This inhibited population increased in prevalence at greater compound concentrations and maintained a saturating rate constant value of constant value of ∼0.0002 s^−1^, consistent with the rate constant observed in high concentrations of Afi by Hartman *et al*. They also found that Afi binds between the upper and lower 50 kDa domains of myosin, same as the blebbistatin binding pocket^12^. This could explain how a similar slowing of ATP turnover occurred in the presence of high concentrations of blebbistatin^36^. This was attributed to the stabilization of switch 2 in myosin heads to limit phosphate release.^38^ Future studies could determine if a similar mechanism occurs with Afi. Overall, these results show that Afi causes greater inhibition of the enzymatic activity of myosin as compared to Mava^36^ without inducing large-scale changes in myosin crown organization or positions relative to the thick filament backbone.

One limitation of the Cy3-ATP experiments was the inability to perform the myofibril biochemical assessments under load. We were not able to pull on the ends of the myofibrils to stretch them to longer sarcomere lengths such as the 2.3 μm sarcomere length that was utilized for the x-ray diffraction and contractility experiments. Few publications have performed biochemical ATP turnover experiments utilizing permeabilized cardiomyocytes. Future work in our group aims to address this limitation and better understand how stretch can impact myofibril nucleotide cycling kinetics.

### Biochemical inhibition does not significantly impact the crossbridge kinetics of myosin heads actively cycling during contraction

Unregulated in-vitro motility results suggest that Afi reduces the number of myosin heads available to interact with actin without incurring significant drag associated with a decrease in fraction moving (Supplemental Figure S3, matching results by Hartman and colleagues^28^). In our steady-state and dynamic contractility experiments, this reduction in available myosin resulted in inhibited force generation. Steady state permeabilized fiber experiments yield no significant differences in cooperative activation (n_H_), myosin cycling kinetics during tension re-stretch (*k*_TR_), or viscous modulus (myosin attachment and detachment frequencies) at the experimental 2.3 µm sarcomere length. Interestingly, myofibril contractile kinetics show that Afi significantly decreases the duration of thin filament deactivation (t_REL,slow_) and accelerated both the slow and fast phases of relaxation of maximally activated myofibrils back to resting baseline. These significant changes in relaxation kinetics are interesting due to the lack of change in myosin cycling kinetics observed in steady state *k*_TR_ and viscoelastic measurements which suggest no significant changes in the load-dependent ADP release rate of myosin heads that are actively cycling and interacting with actin. With Afi inducing significant changes to relaxation kinetics, we hypothesize that the accelerated relaxation kinetics observed in the myofibrils are induced by a reduced pool of actively interacting myosin heads with actin during the transition from high to low calcium solutions.

### Implications of modulating contractility through biochemical and structural inhibition

By biochemical inhibition, Afi decreases the pool of myosin heads that can interact with actin, reducing the capacity of contractility without structurally sequestering heads along the thick filament backbone. Unlike Mava, this mechanistic pathway preserves the recruitment of myosin heads to actin during activation as illustrated by the lack of significant changes in the steady-state fiber viscous modulus dip-frequency, steady state *k*_TR_, the myofibril *k*_ACT_ rate constant values, and the TT_p_ EHT assessment. These differences, however, raise questions regarding how the biochemical and structural modulation of myosin may impact the ability to recruit myosin under varying external conditions. As presented by Ma et al, the structural and functional cardiac relationships to inotropic interventions like Ca^2+^, increased chronotropy, length-dependent activation, and β-adrenergic stimulation, are persevered in the presence of Mava^39^. These significant findings help describe how Mava increased maximum exercise capacity in patients with obstructive HCM^40^ and a subset of non-obstructive HCM patients.^41^ In future studies, it will be important to see how the biochemical inhibition of Afi impacts cardiac reserve, and if the compound can preserve these relationships similar to Mava..

## Conclusion

In conclusion, our study shows that the main mechanism of Afi inhibition is through inhibition of myosin S1 head activity that reduces the number of myosin heads that bind and cycle during contraction. This reduction on recruitable myosin is achieved *without the structural sequestration of the myosin heads on the thick filament backbone*, a significant difference to the mechanistic pathway described for Mava. This finding distinctly decouples the association of biochemical SRX state from the structural ON/OFF states as a mechanism for cardiac muscle inhibition. It also confirms the emergence of an even slower rate of ATP turnover as Afi increases in concentration^28^. Afi does not significantly impact the steady-state cycling kinetics of the myosin heads engaging in contraction and can effectively induce a hypo-contractile phenotype in treated cardiac muscle. The development and success of Afi in the clinic provides a positive outlook for patients suffering with hypertrophic cardiomyopathy along with the field of cardiovascular research.

## Non-standard Abbreviations and Acronyms

Afi: Aficamten
Mava: Mavacamten
HCM: Hypertrophic cardiomyopathy
SL: Sarcomere length
pCa: negative log of calcium concentration
P_i_: inorganic phosphate

## Acknowledgments

The authors acknowledge Mr. Darron Marzolf who provided fresh farm pig hearts. This project used resources from the University of Washington Center for Translational Muscle Research supported by NIH grant P30AR074990. This research used resources of the Advanced Photon Source; a US Department of Energy (DOE) Office of Science User Facility operated for the DOE Office of Science by the Argonne National Laboratory under Contract DE-AC02-06CH11357. BioCAT is supported by NIH grant P30 GM138395.

## Supplemental Methods

### Myofibril single-molecule imaging

#### Myofibril isolation

Porcine left ventricular trabecular myofibril suspensions were prepared as follows: a flash-frozen sample of porcine left ventricular trabeculae was rapidly thawed in chilled Prep buffer (20 mM MOPS, 132 mM NaCl, 5 mM KCl, 4 mM MgCl2, 5 mM EGTA, 10 mM NaN3, 5 mM DTT, protease inhibitor cocktail (A32965; Thermo Scientific), pH 7.1 at RT)^1^ and cut into ∼1 mm thick strips. Sample strips were transferred to 500 µl of Prep buffer in a 2 ml microcentrifuge tube for homogenization using a Tissue Ruptor II (Qiagen) at medium-low speed for 10 seconds twice, with a 1-minute rest on ice between. Homogenized tissue was then permeabilized for 30 minutes at 4 °C, rotating, in Permeabilization buffer (Prep buffer + 1% Triton X-100). Myofibrils were collected by centrifugation at 1000 g for 3 minutes and resuspended in chilled Prep buffer. Washes were repeated 3 times to remove all traces of Triton X-100 and myofibril suspensions were diluted or concentrated where necessary to obtain an OD600 of ∼0.6. Myofibril suspensions were stored at 4 °C and discarded at the end of each day.

#### Microfluidic flow cell preparation

Microfluidic flow cells were constructed by adhering a plasma-cleaned borosilicate coverslip (Menzel Gläser, 24 x 40 mm, 1.5 thickness), coated in 5 µg/ml >300KDa poly-L-lysine (PLL, Sigma) to a standard microscope slide prepared for microfluidics, using a 360 µm thick gasket. Microscope slides were prepared for microfluidics by drilling two 3 mm diameter holes, 1.5 cm apart and adhering polyethylene tubing (Gradko, GE-0086-033, 0.86 mm internal diameter, 0.33 mm wall) to each using an acrylic bonder (RS, 144-406).

### Demembranated Tissue Mechanics Solutions

Frozen porcine left ventricular tissue were sliced into thin strips, thawed, and permeabilized in 50:50 glycerol relaxing solution containing (in mM) 100 KCl, 10 MOPS, 5 K_2_EGTA, 9 MgCl_2_ and 5 Na_2_ATP (adjusted to pH = 7 with KOH), 1% (by vol) Triton X-100, 1% protease inhibitor (sigma P8340), and 50% (by vol) glycerol at 4°C overnight. The solution was changed to the same 50:50 glycerol relaxing solution without Triton X-100 for storage up to one week at −20°C.

pCa solutions were calculated using an in-house Excel workbook. Ionic strength was set to 170 mM and pH to 7.0 at 21°C. Total component concentrations included (in mM) 15 EGTA, 80 MOPS, 5 MgATP, 15 creatine phosphate, 83 free K, and 52 free Na. Calcium was varied from pCa 9.0 to pCa 4.5. pH was adjusted, maintaining ionic strength using HCl and TrisOH.

### In vitro motility assay

Myosin and heavy meromyosin (HMM) were purified from pig left ventricular samples as previously described.^2^ *In vitro* motility assays were performed at 30 °C using unregulated Rhodamine Phalloidin labeled F-actin in the presence of 2 mM ATP and either DMSO, 1 μM Afi, or 1 μM Mava.^2,3^ The concentration of HMM was varied from 0.2 mg/ml down to 0.04 mg/ml and moving velocity and fraction moving were assessed. Custom-built software analyzed images of the moving filaments as in our previous publications.^2,3^

### Human induced pluripotent stem cell derived cardiomyocyte (hiPSC-CM) differentiation

Undifferentiated human induced pluripotent stem cells (hiPSCs) (WTC11) were thawed from liquid nitrogen storage into a solution of mTeSR-1 (StemCell Technologies) + 10μM Y-27632 ROCK inhibitor. These cells were maintained on tissue culture plastic coated with Matrigel (Corning) diluted to a concentration of 5.1 mg/mL, at 37°C and 5% CO2. 24 hours later, this media was replaced with fresh mTeSR. These cells were maintained for a 3 days post-thaw, before dissociation with Accutase (StemCell Technologies) replating to control plate confluency. 3 days post replate, cells were plated in a 12 well plate at 0.5-1.5×10^5 cells/well. 24 hours after plating, media was replaced with mTeSR + 1 μM CHIR99021 (Chiron). 48 hours after plating, differentiation was initiated by changing the media to 4-5 μM Chiron (Caymen chemical) in RPMI 1640 + Bovine Serum Albumin + Ascorbic Acid (RBA media). 48 hours post-induction, the media was changed to RBA supplemented with WNT-C59. 48 hours after WNT inhibition, media was changed to fresh RBA. Finally, 48 hours after the removal of WNT-C59, media was switched to 2mL/well of RPMI + B27+insulin and changed every 48 hours. Spontaneous contraction was observed on day 6 post-induction.

## Supplemental Tables

**Table S1.**
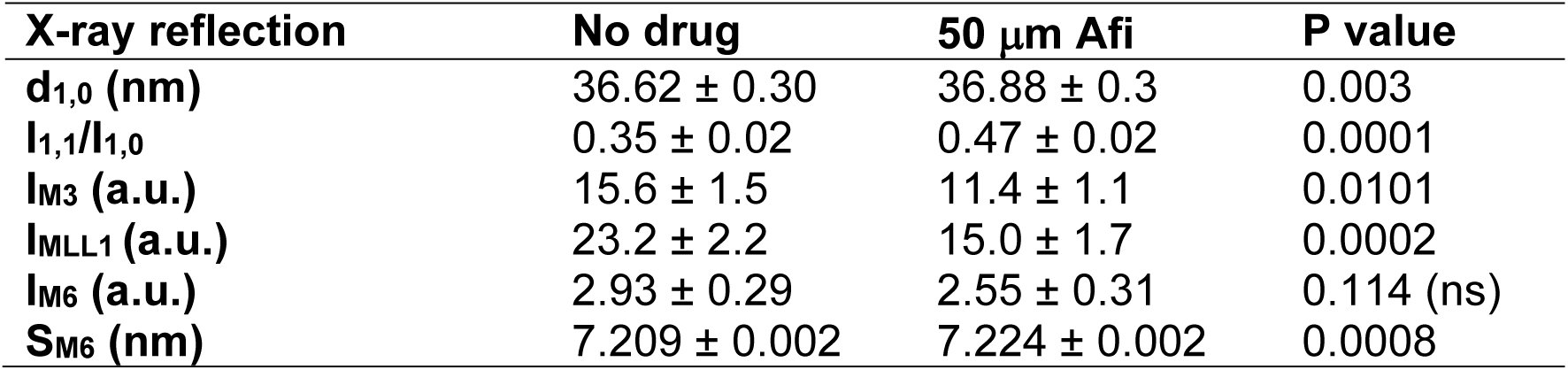
Numerical values of X-ray diffraction reflections of relaxing muscle (pCa 8.0). Values represent mean ± S.E.M. for n= X preparations using paired two-tailed t-test analysis.

| **X-ray reflection** | **No drug** | **50 μm Afi** | **P value** |
| --- | --- | --- | --- |
| **d<sub>1,0</sub> (nm)** | 36.62 ± 0.30 | 36.88 ± 0.3 | 0.003 |
| **I<sub>1,1</sub>/I<sub>1,0</sub>** | 0.35 ± 0.02 | 0.47 ± 0.02 | 0.0001 |
| **I<sub>M3</sub> (a.u.)** | 15.6 ± 1.5 | 11.4 ± 1.1 | 0.0101 |
| **I<sub>MLL1</sub> (a.u.)** | 23.2 ± 2.2 | 15.0 ± 1.7 | 0.0002 |
| **I<sub>M6</sub> (a.u.)** | 2.93 ± 0.29 | 2.55 ± 0.31 | 0.114 (ns) |
| **S<sub>M6</sub> (nm)** | 7.209 ± 0.002 | 7.224 ± 0.002 | 0.0008 |

## Supplemental Figures

**Figure S1.**
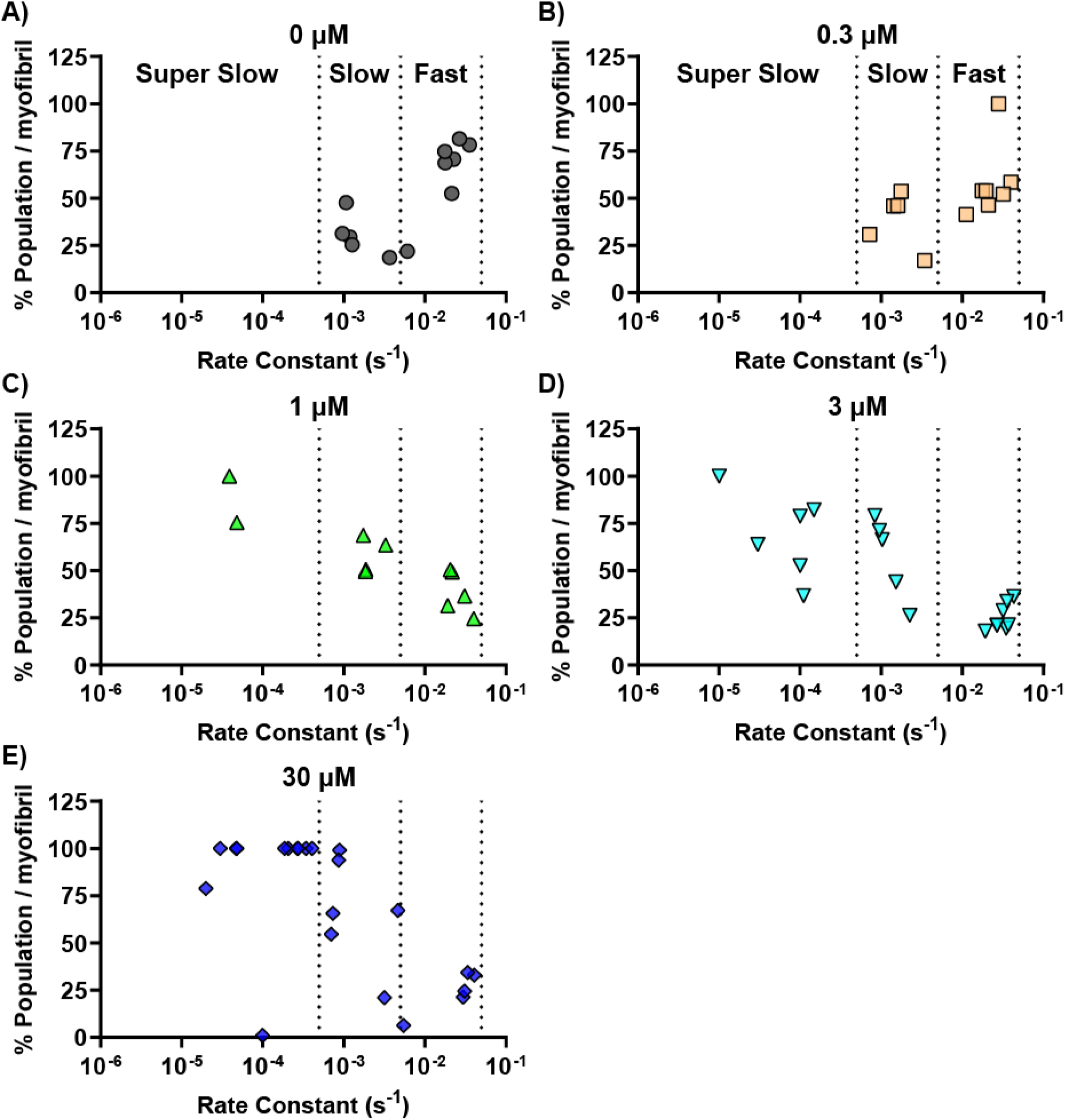
Increasing concentrations of Afi moves the percentage of rate constant populations for each myofibril away from fast state, into slow state, and eventually into the super slow state that emerges at concentrations at 1 μΜ and higher. Each rate constant within a myofibril is normalized based on the relative amplitudes so that the total rate constant population is 100% for each myofibril. Each rate constant is plotted as a single point and binned into fast (0.05 - 0.005 s^−1^), slow (0.005 - 0.0005 s^−1^), or super slow (< 0.0005 s^−1^) states at concentrations of Afi including A) 0 μM, B) 0.3 μM, C) 1 μM, D) 3 μM, and E) 30 μM.

**Figure S2.**
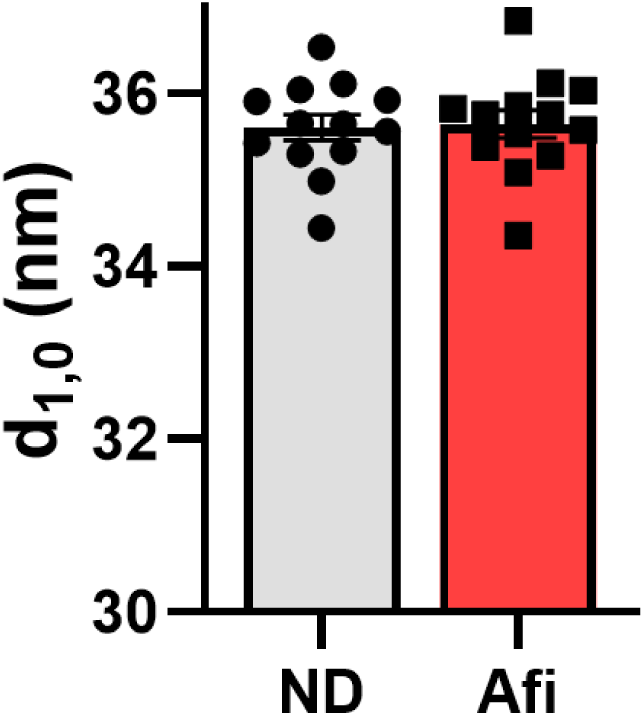
Incubation in 50 μM Afi does not alter cardiac lattice spacing (d_1,0_).

**Figure S3.**
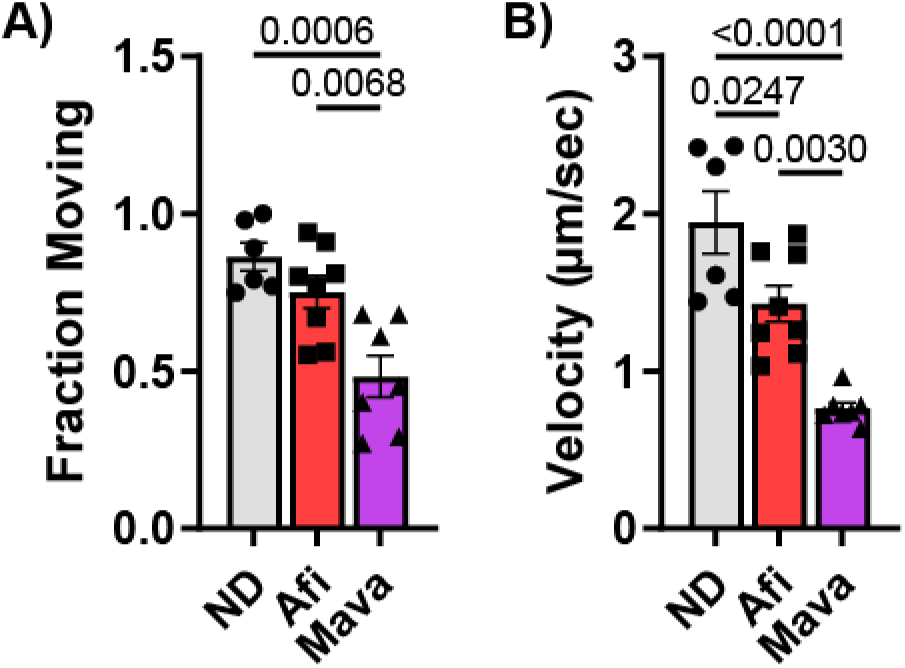
Afi and Mava decrease filament velocity. A) Only 1 μM Mava significantly decreases the fraction of actin filaments moving in comparison to ND, while 1 μM Afi does not. B) Both Afi and Mava significantly decrease actin sliding velocity, with Mava to a greater extent. N = 6-8 slides per condition with statistics calculated using one-way ANOVA with Tukey’s multiple comparison test.

